# Monitoring migratory birds in stopover habitat: assessing the value of extended duration audio recording

**DOI:** 10.1101/2020.08.01.232215

**Authors:** Ellie Roark, Willson Gaul

## Abstract

1. Because birds are frequently detected by sound, autonomous audio recorders (called automated recording units or ARUs) are now an established tool in addition to in-person observations for monitoring the status and trends of bird populations. ARUs have been evaluated and applied during breeding seasons, and to monitor the nocturnal flight calls of migrating birds. However, birds behave differently during migration stopover than during the breeding season. Here we present a method for using ARUs to monitor land birds in migration stopover habitat.
2. We conducted in-person point counts next to continuously recording ARUs, and compared estimates of the number of species detected and focal species relative abundance from point counts and ARUs. We used a desk-based audio bird survey method for processing audio recordings, which does not require automated species identification algorithms. We tested two methods of using extended duration ARU recording: surveying consecutive minutes, and surveying randomly selected minutes.
3. Desk-based surveys using randomly selected minutes from extended duration ARU recordings performed similarly to point counts, and better than desk-based surveys using consecutive minutes from ARU recordings. Surveying randomly selected minutes from ARUs provided estimates of relative abundance that were strongly correlated with estimates from point counts, and successfully showed the increase in abundance associated with migration timing. Randomly selected minutes also provided estimates of the number of species present that were comparable to estimates from point counts.
4. ARUs are an effective way to track migration timing and intensity in remote or seasonally inaccessible migration stopover habitats. We recommend that desk-based surveys use randomly sampled minutes from extended duration ARU recordings, rather than using consecutive minutes from recordings. Our methods can be immediately applied by researchers with the skills to conduct point counts, with no additional expertise necessary in automated species identification algorithms.

## 1 INTRODUCTION

Conserving bird populations requires knowledge of bird distribution and habitat use at all stages of their life cycle, including during breeding, migration, and non-breeding periods (Sherry & Holmes, 1995). Monitoring birds’ habitat use during migration is a necessary component of conservation plans for migratory birds. Historically, researchers have primarily relied on in-person observations including mist-netting (Peach, Buckland & Baillie, 1996) and point counts (Ralph, Droege & Sauer, 1995) for migration monitoring, but because birds are frequently detected by sound, audio recording technology offers opportunities to expand monitoring techniques. Here we present a method for using audio recorders to monitor birds in migration stopover habitat during spring migration.

Figuring out how to best monitor bird abundance and diversity in remote habitat is a current challenge. The climate in high latitude continental regions increases the challenges associated with accessing remote areas during spring migration. Significant annual snow accumulation, followed by rapid melting as temperature increases, makes unpaved roads impassable for several weeks each spring in much of northern North America, typically during the same time period when migrant bird species begin to arrive in the region. Developing survey monitoring protocols that can be implemented despite poor traveling conditions is a way to fill in gaps in knowledge of northern forest birds, and birds in similarly remote habitats.

Autonomous recording units (ARUs) are programmable audio recorders that can be deployed in the field for long time periods to efficiently maximize the spatial and temporal extent of monitoring. Passive acoustic monitoring is widely used in ecology to monitor and study vocalizing organisms; ARUs have been deployed to study bats (Tuneu-Corral et al., 2020), whales (Baumgartner et al., 2019), invertebrates (Penone et al., 2013), amphibians (Dutilleux & Curé, 2020) and birds (Shonfield & Bayne, 2017). ARUs are also used to evaluate the success of conservation programs (Shonfield & Bayne, 2017). Current challenges for implementing passive acoustic monitoring include the availability of reference sound libraries, minimizing errors in species identification, and determining the relationship between acoustic index values and their associated real-world underlying parameters (Gibb, Browning, Glover-Kapfer & Jones, 2019).

Point-count surveys are the most commonly used bird monitoring protocol for long-term study sites (Ralph, Droege, & Sauer, 1995; Rosenstock, Anderson, Giesen, Leukering & Carter, 2002), but ARUs are now viewed as a viable supplement to point-counts, especially during the breeding season when birds vocalize frequently (Furnas & Callas, 2015; Klingbeil & Willig, 2015; Shonfield & Bayne, 2017; Darras et al., 2018; Darras et al., 2019). Many researchers have compared ARUs and point counts in terms of their estimates of species richness and relative abundance or occupancy (Haselmayer & Quinn, 2000; Campbell & Francis, 2011; Tegeler, Morrison & Szewczak, 2012; La & Nudds, 2016), including in temperate forest (Klingbeil & Willig, 2015). However, none of these studies (including the 23 studies reviewed in Darras et al.’s (2018) meta-analysis) compared point counts and ARUs during migration. Birds behave and vocalize differently during migration than during the breeding season (Morse, 1991; Rappole & Warner, 1976). Testing and refining migration-specific monitoring techniques for ARUs is therefore necessary to understand how data from ARUs compare to data from in-person observations.

ARUs are currently used during migration to record the flight calls of nocturnally migrating species. They are deployed to track the abundance of migrants as they move through an area, and can provide helpful information about migratory flyway locations, migration phenology, and relative abundance (Sanders & Mennill, 2014; Evans & Rosenburg, 2000). Understanding how migrating birds use migratory stopover habitat is a different challenge, and requires different methods. Determining how birds are distributed in stopover habitat, the relative abundance and species richness of birds in such habitat, and the timing of arrival and departure from the stopover area are all important research questions for applied conservation.

To take advantage of the large volume of data generated by continuously recording ARUs, researchers are actively developing methods for automated identification of vocalizing organisms (Salamon et al., 2016; Gibb, Browning, Glover-Kapfer & Jones, 2019). In contrast, we present a method that can be implemented by anyone with the skills to conduct point counts, that does not rely on machine learning for species identification and data processing. Because applications of ARUs in migration stopover habitat have been under-explored in the literature thus far, we demonstrated and assessed an immediately applicable monitoring technique.

We compared data from ARU surveys to in-person point count surveys during spring migration in the northern Great Lakes region of the United States. Our goal was to understand how ARUs could be applied to monitor diurnal stopover habitat use during migration by examining whether ARUs could provide estimates of relative abundance and number of species that are comparable to estimates from in-person surveys. We asked the following questions. 1) What are the differences between the number of species detected using point counts and using ARUs? 2) Can ARUs give estimates of relative abundance for focal species that are correlated with estimates of relative abundance from point counts? 3) Can randomly sampling from extended duration audio recordings provide better estimates of focal species abundance or the number of species detected than consecutive minutes of audio recording?

## 2 MATERIALS AND METHODS

We conducted in-person point counts alongside continuously recording ARUs on the southern shore of Lake Superior during two months at the start of spring migration. We compared both raw data and model-based estimates of the number of species detected and focal species abundance from point counts and ARUs.

### 2.1 Study site

We conducted field work in a 2.7 km^2^ area on the Point Abbaye peninsula in Baraga County, Michigan, USA (Fig. 1). Surveys took place from 2 April to 22 May 2019, and were conducted daily unless prevented by weather conditions. Field work was designed to coincide with the arrival and peak abundance of early season migrating birds. Point Abbaye juts into the southern part of Lake Superior and comprises the western border of Keweenaw Bay. Habitat included forested wetland, upland hardwood, and hardwood forest disturbed by recent logging activity. We selected survey sites randomly across the study area. All spatial analyses were done in the R programming language using the ‘rgdal’, ‘geosphere’, ‘rgeos’, ‘sp’, ‘maptools’, and ‘spatstat’ packages (Baddeley, Rubak, & Turner, 2015; Bivand, Keitt, & Rowlingson, 2018; Bivand & Lewin-Koh, 2019; Bivand & Rundel, 2018; Bivand, Pebesma, & Gomez-Rubio, 2013; Hijmans, 2019; Pebesma & Bivand, 2005; R Core Team, 2020). We conducted a pilot study in 2018 to test our protocols and evaluate the accessibility of our randomly selected survey locations. See Appendix A for details about pilot year surveys, and survey site and date selection.

**Figure 1:**
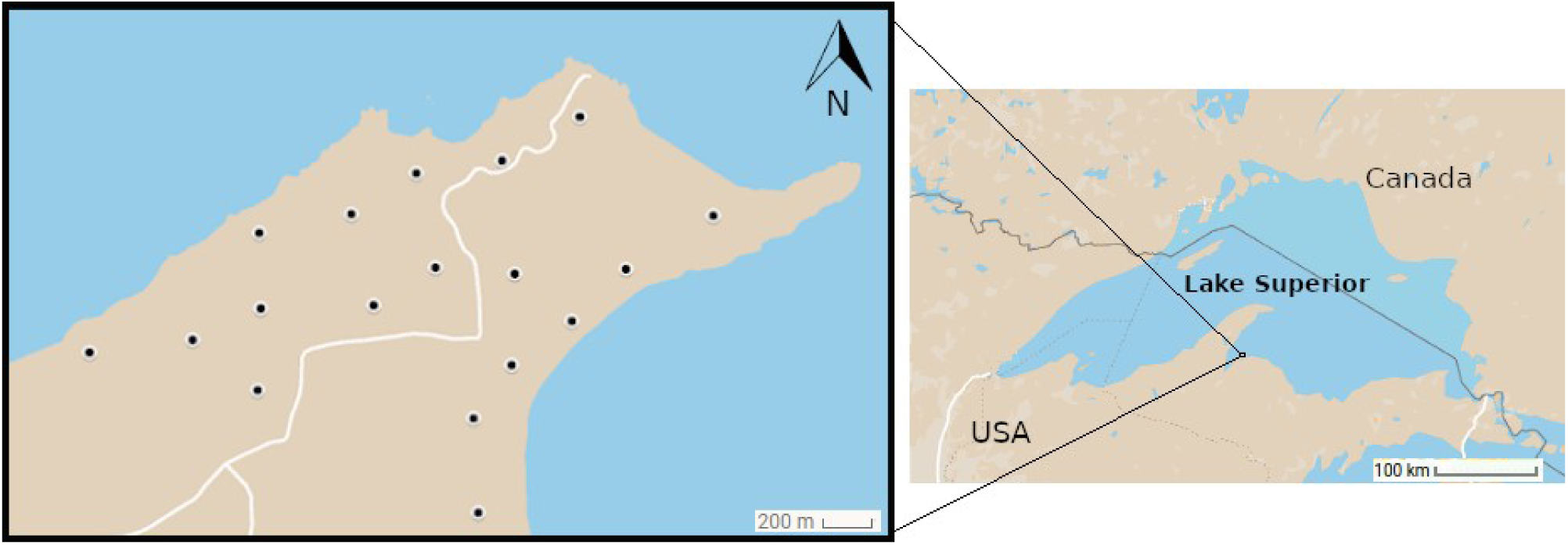
Study area on the Point Abbaye Peninsula in Baraga County, Michigan, USA. Points denote survey locations visited during April and May of 2019. The survey area covers 2.7 km^2^. White lines on the left panel show unploughed four wheel drive roads which are impassible from early April to early May each year.

### 2.2 Automated recording units

Birds were recorded using three SWIFT bioacoustic recorder rugged units (Cornell Lab of Ornithology, Ithaca, NY, USA), and one AudioMoth bioacoustic recorder that was housed in a thin plastic bag for light weather proofing (Hill et al., 2018; Open Acoustic Devices, Southampton, UK). SWIFT units used a built in PUI Audio brand omni-directional microphone. The AudioMoth unit used an analog microelectro-mechanical systems (MEMS) microphone. We refer to both the SWIFT and AudioMoth units as “automated recording units” (ARUs). ARUs recorded at a sampling rate of 48 kHz and saved recordings as uncompressed .WAV files. The signal to noise ratio reported by device manufacturers is approximately 58 dB for the SWIFT units, and approximately 44 dB for the AudioMoth unit.

### 2.4 Field survey methods

ARUs recorded continuously for five hours each day, beginning within 10 minutes of local sunrise time (United States Naval Observatory, 2016). ARUs were attached to trees less than 0.6 m in diameter, and were placed 1.5–2 m above the ground (Darras et al., 2018). The SWIFT omni-directional microphones were always oriented downward to prevent precipitation landing directly on the microphone. After the five hour recording period ended each day, ARUs were moved to new locations for the next day’s samples, thereby rotating the ARU and point count samples through all 18 survey locations approximately every five days. The sampling order for the points was chosen randomly.

Point counts were conducted daily next to each ARU during the five hour recording period. Point counts involved recording all birds seen and heard at an unlimited distance during a stationary, 10-minute count. We did not survey in high wind or heavy precipitation. See Appendix A for detailed point count protocols.

### 2.5 Desk-based audio surveys

We conducted desk-based audio bird surveys by listening to ARU recordings played through headphones on a laptop computer in the lab after the end of the field season. We tested three types of desk-based audio surveys: 1) we listened to a recording of the 10 consecutive minutes during which the in-person point count was conducted; 2) we listened to 24 minutes selected randomly from the five hour recording duration; 3) we sampled a subset of 10 of the 24 random minutes (without listening to those minutes again). Our goal was to compare each of these desk-based ARU survey methods to in-person point count observations.

For each audio file, the desk-based survey technician noted the identity of each bird species that vocalized, the type of vocalization, and the 30-second time intervals in which each species vocalized. Detailed protocols for completing desk-based audio surveys can be found in Appendix A, and a completed data sheet from a desk-based survey is shown in Fig. S6. We sampled 24 random minutes from each ARU on each day because we wanted at least 20 minutes without anthropogenic disturbance. After discarding randomly selected minutes that contained a human voice, we were ultimately able to use data from 22 randomly selected minutes from each ARU on each survey day (i.e. we never had to discard more than two of the 24 randomly selected minutes because of human voices).

### 2.6 Indices for observed number of species and relative abundance

We summed the number of unique species detected (*S*) separately using each survey type: 10-minute, in-person point counts (*S*_p_); 10 consecutive minute ARU surveys (*S*_10C_); 10 random minute ARU surveys (*S*_10R_); and 22 random minute ARU surveys (*S*_22R_) (Box 1). We calculated a value of *S* for each individual survey on each day, resulting in three or four values of *S* for each survey type on each day.

#### Box 1

Indices of relative abundance and species richness for both in-person point count observations, and desk-based listening counts using audio data from Automated Recording Units (ARUs).

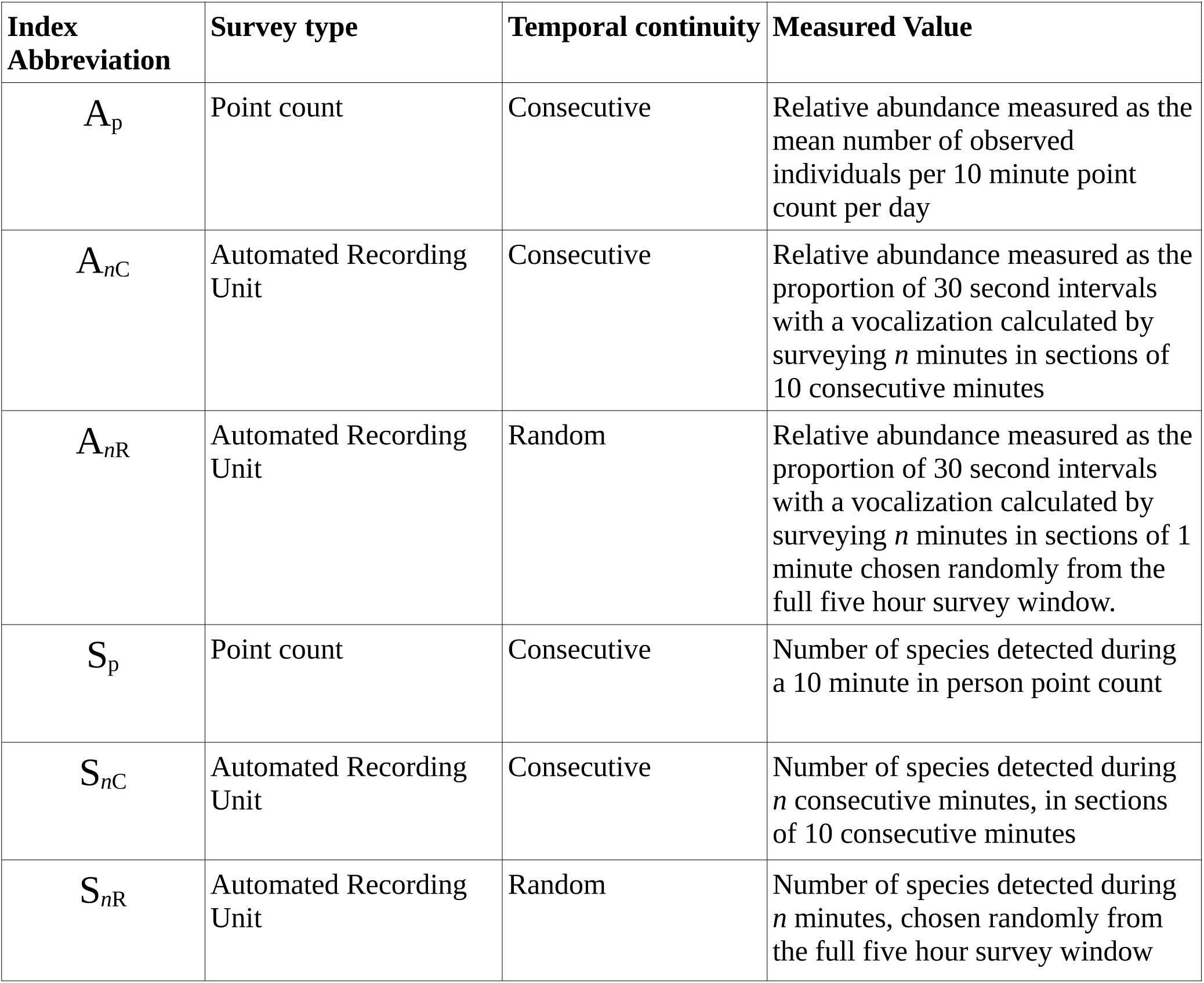

We created an index of daily relative abundance (*A*) for two focal species (*Regulus satrapa* (Golden-crowned Kinglet) and *Troglodytes hiemalis* (Winter Wren)) using each survey type (Box 1). Our relative abundance indices were: the mean observed number of individuals per point count (*A*_p_); the proportion of 30-second intervals with a vocalization calculated by surveying *n* minutes in sections of 10 consecutive minutes (*A*_*n*C_); the proportion of 30-second intervals with a vocalization calculated by surveying *n* minutes in sections of one minute chosen randomly from the five hour survey window (*A*_*n*R_). To reduce the number of zero abundance counts in our data, we calculated relative abundance indices by grouping all surveys of each type for each day, so there was a single value for each abundance index on each day. Note that while April 16^th^ and 17^th^ data appear on plots and in results, ARU malfunctions on those dates made the number of sampled minutes *n* different for those two dates for some of our abundance indices. See Appendix A for detailed discussion of sample size on these dates.

### 2.7 Statistical Analysis

#### 2.7.1 Observed number of species

To determine whether the survey type (*S*_10C_, *S*_10R_, *S*_22R_) significantly influenced the number of species detected, we modeled the number of species detected using a generalized linear mixed model (GLMM) with a Poisson error distribution and log link function, using the ‘lme4’ package in R (Bates, Maechler, Bolker, & Walker, 2015; R Core Team, 2020). Our fixed effects were survey type, day of year, a second degree polynomial term for day of year, wind, rain, noise, and interaction terms for day of year x survey type, and survey type x rain. We also used day of year as a random effect; we collected up to four samples of each of four survey types per day, and new birds potentially arrived daily during the study period, so we expected that the number of species detected by all surveys on each day would be strongly correlated, regardless of survey location or survey type. More information about our GLMM can be found in Appendix A.

#### 2.7.2 Relative abundance

We compared relative abundance estimates for our two focal species using data from *A*_p_ and from each of the three desk-based audio survey types (*A*_30C_, *A*_30R_, *A*_66R_). Winter Wrens were abundant in the survey area, and vocalized frequently and loudly during early spring, representing a “best case” scenario for detectability on ARU recordings. Golden-crowned Kinglets were abundant in the survey area, but vocalized quietly (though regularly) during early spring, and so represent a greater challenge for detection using ARUs.

We produced a total of eight relative abundance models, one using each of our four relative abundance indices for each of our two focal species. For each relative abundance model, we fit boosted regression trees (BRTs) (Elith, Leathwick, & Hastie, 2008; Friedman, 2001) using 200 iterations of five-fold temporal block cross validation, that used blocks of three consecutive days (Fig. S3; Roberts et al., 2017). This resulted in a total of 1000 BRT fits per relative abundance model. We generated predicted relative abundances by averaging predictions from the 1000 fits of each model. The predictor variables in our model were day of year (continuous) and wind speed (categorical with three levels representing Beaufort forces of 0-1, 2, or 3 or higher). We fit BRTs with the ‘gbm’ package in R (Greenwell, Boehmke, Cunningham and GBM Developers, 2019; R Core Team, 2020). Details of BRTs, including model tuning and control of overfitting are in Appendix A, and Figs. S4-S5.

We assessed the correlation between the observed data from our four abundance indices using scatter plots and Spearman’s rank correlation coefficient. We also assessed correlation between the predicted values from the BRT models trained using data from each of the four abundance indices. We calculated Spearman’s rank correlation coefficient for all pairwise combinations of abundance indices, with the exception A_30R_ and A_66R_, since the data for A_30R_ is a sub-sample of A_66R_ data.

## 3 RESULTS

Between 2 April and 22 May, 2019, we were able to survey on 37 days. During that time, we conducted 137 in-person point counts. All four audio recorders experienced occasional malfunctions that prevented us from recording the full five-hour survey window with some units on some days. One SWIFT unit recorded 36 five-hour survey days, two SWIFT units recorded 35 five-hour survey days, and the AudioMoth unit recorded 24 five-hour survey days, for a total of 650 hours recorded by ARUs. Because of ARU malfunctions, on some days ARUs did not record the full five-hour period, but we were able to manually turn on the units for the 10-minute period during the in-person point count.

Therefore, we recorded 130 10 consecutive minute periods with ARUs (during which a human observer was present conducting a simultaneous point count) on 37 survey days, but only 124 periods of 22 randomly selected minutes on 36 survey days. A complete list of the species detected by each survey method can be found in Table S1.

### 3.1 Observed number of species

A chi-square ANOVA comparing our full model to a null model with survey type removed showed that survey type (the *S*-index used) had a significant effect on the number of species detected (χ^2^ _9, 21_=247, *p* < 0.0001). We detected a similar number of species using *S*_10R_ as we did using *S*_p_ (Fig. 2; Table 1; change in the log of the number of species detected = −0.065, 95% CI [-0.2; 0.08], *p =* 0.3888). Using *S*_22R_, we detected significantly more species than by using *S*_p_ (Fig. 2; Table 1; change in the log of the number of species detected = 0.305, 95% CI [0.17; 0.44], *p* < 0.0001). We detected fewer species using *S*_10C_ than using *S*_p_ (Fig. 2; Table 1; change in the log of the number of species detected = −0.614, 95% CI [−0.78; −0.44], *p* < 0.0001). Listening to randomly selected rather than consecutive minutes eliminated the gap in number of species detected between 10-minute point counts and 10-minute ARU surveys (Fig. 2). Day of year had a significant effect on the number of species detected (Table 1), with more species expected later in the migration season (Fig. 2).

**Table 1:** Results of a generalized linear mixed model of the number of species detected as a function of day of year, count type and environmental condition covariates. Each variable included in the model is shown, along with the coefficient point estimate, 95% confidence interval, and significance level. *denotes p <0.05, ** denotes p <0.01, *** denotes p < 0.001

| Variable | Coefficient estimate | 95% CI lower bound | 95% CI upper bound | P-value |
| --- | --- | --- | --- | --- |
| Survey type (S<sub>10C</sub>) | -0.6146 | -0.7824 | -0.4467 | <0.0001*** |
| Survey type (S<sub>10R</sub>) | -0.065 | -0.213 | 0.0829 | 0.3888 |
| Survey type (S<sub>22R</sub>) | 0.3056 | 0.1702 | 0.441 | <0.0001*** |
| Wind (2) | -0.0301 | -0.135 | 0.0748 | 0.574 |
| Wind (3+) | -0.1007 | -0.2644 | 0.0629 | 0.2277 |
| Rain (Wet) | -0.1984 | -0.4624 | 0.0657 | 0.1409 |
| Noise (1) | 0.1552 | 0.0595 | 0.251 | 0.0015** |
| Noise (>2) | 0.0544 | -0.147 | 0.2557 | 0.5967 |
| Day of year | 0.4346 | 0.305 | 0.5641 | <0.0001*** |
| Day of year<sup>2</sup> | -0.1159 | -0.2496 | 0.0178 | 0.0892 |
| Survey type (S<sub>10C</sub>) x Rain (Wet) | -0.0163 | -0.3944 | 0.3617 | 0.9325 |
| Survey type (S<sub>10R</sub>) x Rain (Wet) | 0.1861 | -0.1374 | 0.5096 | 0.2595 |
| Survey type (S<sub>22R</sub>) x Rain (Wet) | 0.2288 | -0.0677 | 0.5252 | 0.1304 |
| Survey type (S<sub>10C</sub>) x Day of Year | 0.0153 | -0.1191 | 0.1498 | 0.823 |
| Survey type (S<sub>10R</sub>) x Day of Year | 0.0595 | -0.0659 | 0.1849 | 0.3525 |
| Survey type (S<sub>22R</sub>) x Day of Year | -0.0709 | -0.1808 | 0.0389 | 0.2058 |
| Survey type (S<sub>10C</sub>) x Day of year<sup>2</sup> | 0.086 | -0.048 | 0.2199 | 0.2083 |
| Survey type (S<sub>10R</sub>) x Day of year<sup>2</sup> | -0.0583 | -0.1825 | 0.0659 | 0.3575 |
| Survey type (S<sub>22R</sub>) x Day of year<sup>2</sup> | -0.0148 | -0.125 | 0.0953 | 0.7917 |

**Figure 2:**
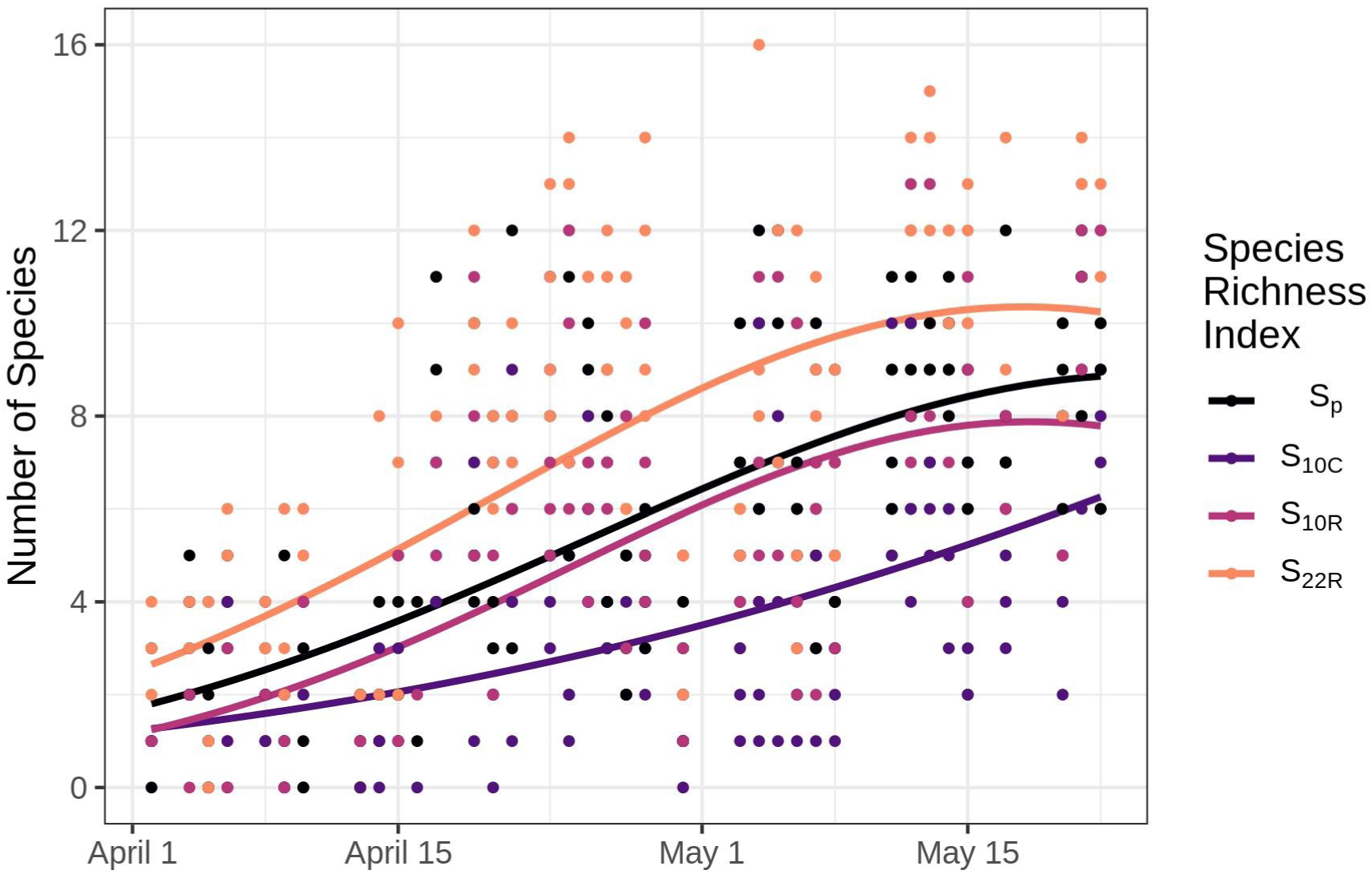
The number of bird species detected by in-person point counts and automated recording units (ARUs) during the spring migration period on the Point Abbaye peninsula, Michigan, USA in 2019. Points show observed number of species detected, lines show predictions from a generalized linear mixed model, holding weather variables constant. See Box 1 for a description of the species richness index abbreviations. Listening to 10 random minutes of data from an ARU (*S*_10R_) allowed for detection of the same number of species as a 10 consecutive minute in-person point count (*S*_p_). Increasing survey effort to 22 random minutes of ARU data (*S*_22R_) increased the number of species detected to above the number of species detected by in-person point counts.

Chi-square ANOVA showed that the overall effect of wind was not significant (χ^2^ _19, 21_=1.44, *p* = 0.48), nor was the overall effect of rain (χ^2^ _18, 21_= 3.45, *p* = 0.32). The overall effect of noise was significant (χ^2^ _19, 21_= 10.3, *p =* 0.005). The interaction between survey type and day of year was not significant (Table 1), providing no evidence of a difference in the effect of survey method on the observed number of species over the course of the survey season.

### 3.2 Relative abundance models

BRT models of relative abundance over time differed in how well they showed the initial period of absence, and the increase in abundance corresponding with the arrival of migrant birds in our study area, depending on the survey method used (Fig. 3). The general pattern of initial absence followed by arrival of migrants can be seen in both the raw data and the model predictions of relative abundance for *A*_p_, *A*_30R_ and *A*_66R_ for Winter Wrens (Fig. 3 a, c and d), and for *A*_p_ and *A*_66R_ for Golden-crowned Kinglets (Fig. 3 e, and h).

**Figure 3:**
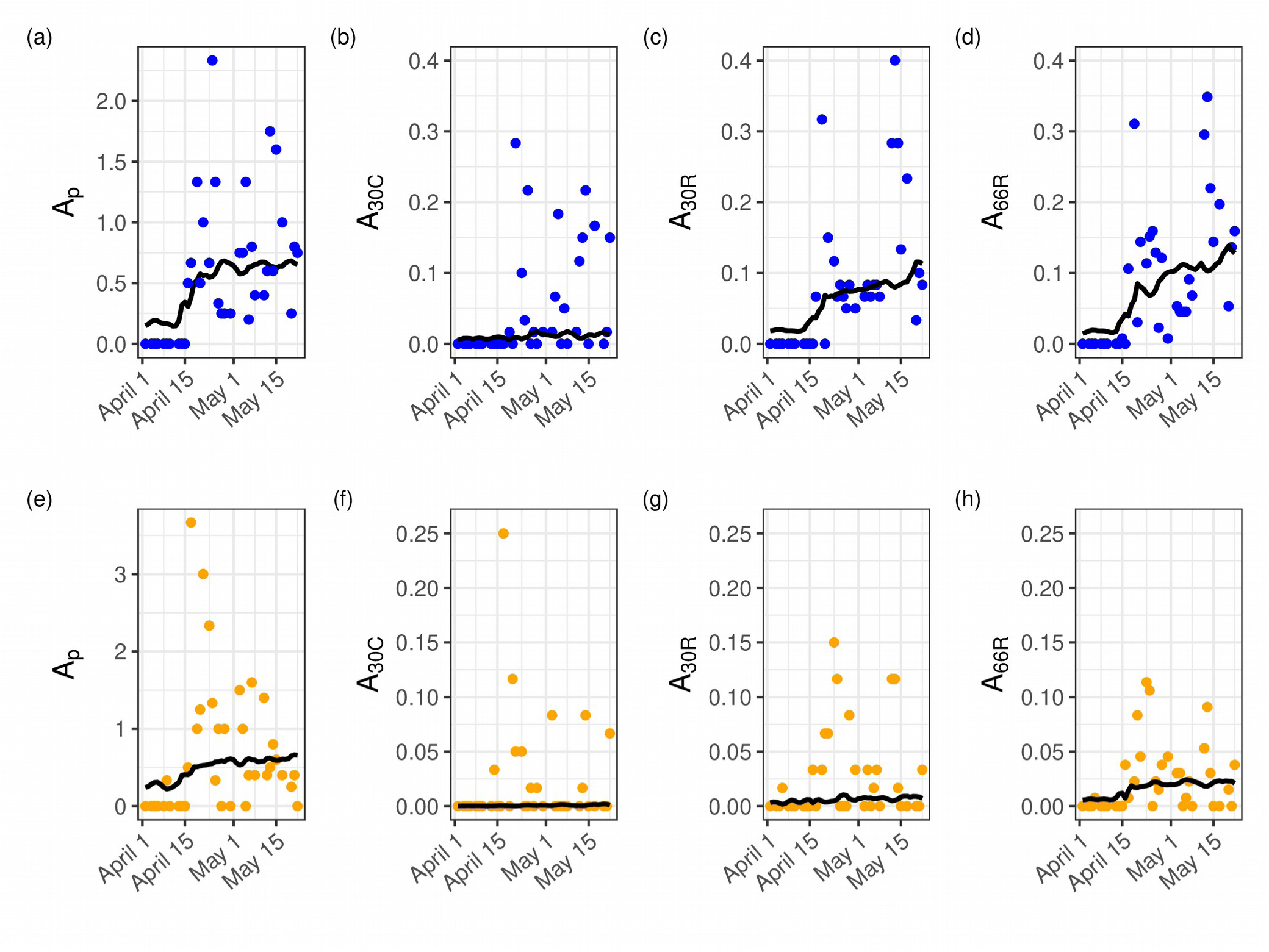
Predicted (lines) and observed (points) relative abundance of (a) through (d) Winter Wren, and (e) through (h) Golden-crowned Kinglet, in April and May of 2019 on the Point Abbaye peninsula. Points show observed values for each abundance index, while lines show the mean predicted values from 200 five-fold cross validated boosted regression tree models. The vertical axes show the daily abundance index value (see Box 1) calculated from: (a) and (e) three 10-minute in person point counts, (b) and (f) three samples of 10 consecutive minutes of audio recordings from automated recording units (ARUs), (c) and (g) three samples of 10 randomly selected minutes of audio recordings from ARUs, (d) and (h) three samples of 22 randomly selected minutes of audio recordings from ARUs.

For both Winter Wrens and Golden-crowned Kinglets, the observed abundance indices from ARU surveys were positively correlated with the observed abundance index from point counts (Fig. 4, Fig. 5), indicating that the relative abundance proxies we calculated using ARUs are comparable to relative abundance estimates from in-person observations. Winter Wren showed moderate to strong correlation between the observed abundance index values from point counts and from ARU surveys (Fig. 4, Table 2). Abundance indices for Golden-crowned Kinglets were less correlated than for Winter Wrens, with weak correlation between *A*_30C_ and *A*_30R_ in particular (Fig. 5, Table 2). Correlations for predicted values of our abundance indices were moderate to strong for both species (Figs. S1-S2), and were higher than correlation coefficients for observed values of the same index pairs (Table 2). The strong correlation between predicted values of the abundance indices indicates that our models found the same underlying signal regardless of whether training data were from ARUs or point counts.

**Table 2:** Spearman’s rank correlation coefficients for each combination of abundance indices. We did not report a correlation for A_30R_ and A_66R_ because A_30R_ data is a subset of A_66R_ data.

| Relative Abundance Index Pairs |  | Golden-crowned Kinglet |  | Winter Wren |  |
| --- | --- | --- | --- | --- | --- |
|  |  | *Correlation of Observed Values* | *Correlation of Predicted Values* | *Correlation of Observed Values* | *Correlation of Predicted Values* |
| A<sub>P</sub> | A<sub>30C</sub> | 0.532 | 0.794 | 0.72 | 0.822 |
| A<sub>P</sub> | A<sub>30R</sub> | 0.499 | 0.652 | 0.781 | 0.882 |
| A<sub>P</sub> | A<sub>66R</sub> | 0.588 | 0.813 | 0.817 | 0.854 |
| A<sub>30C</sub> | A<sub>30R</sub> | 0.286 | 0.817 | 0.716 | 0.757 |
| A<sub>30C</sub> | A<sub>66R</sub> | 0.485 | 0.897 | 0.676 | 0.736 |

**Figure 4:**
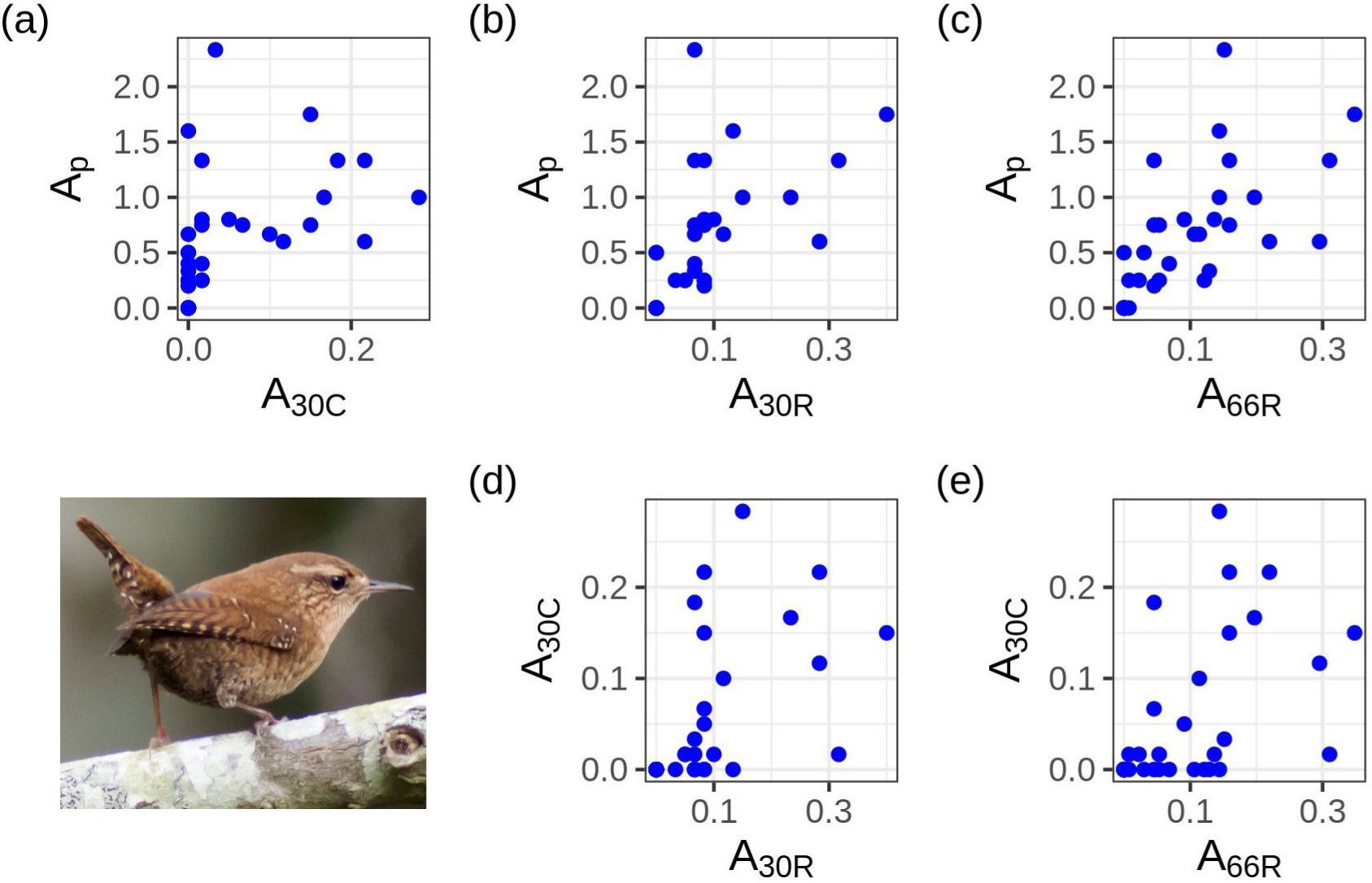
Correlation between daily observed values of relative abundance indices (Box 1) for Winter Wren. See Table 2 for Spearman’s correlation coefficients for each plot. Abundance indices for ARUs (the proportion of 30-second intervals with a vocalization) are correlated with the abundance index from point counts (mean number of individuals observed per count per day). Note that axis scales vary by abundance index; absolute values are less important here than the relationship between observations. Photo: “Winter Wren” by ilouque, used under license CC BY 2.0. Cropped from original.

**Figure 5:**
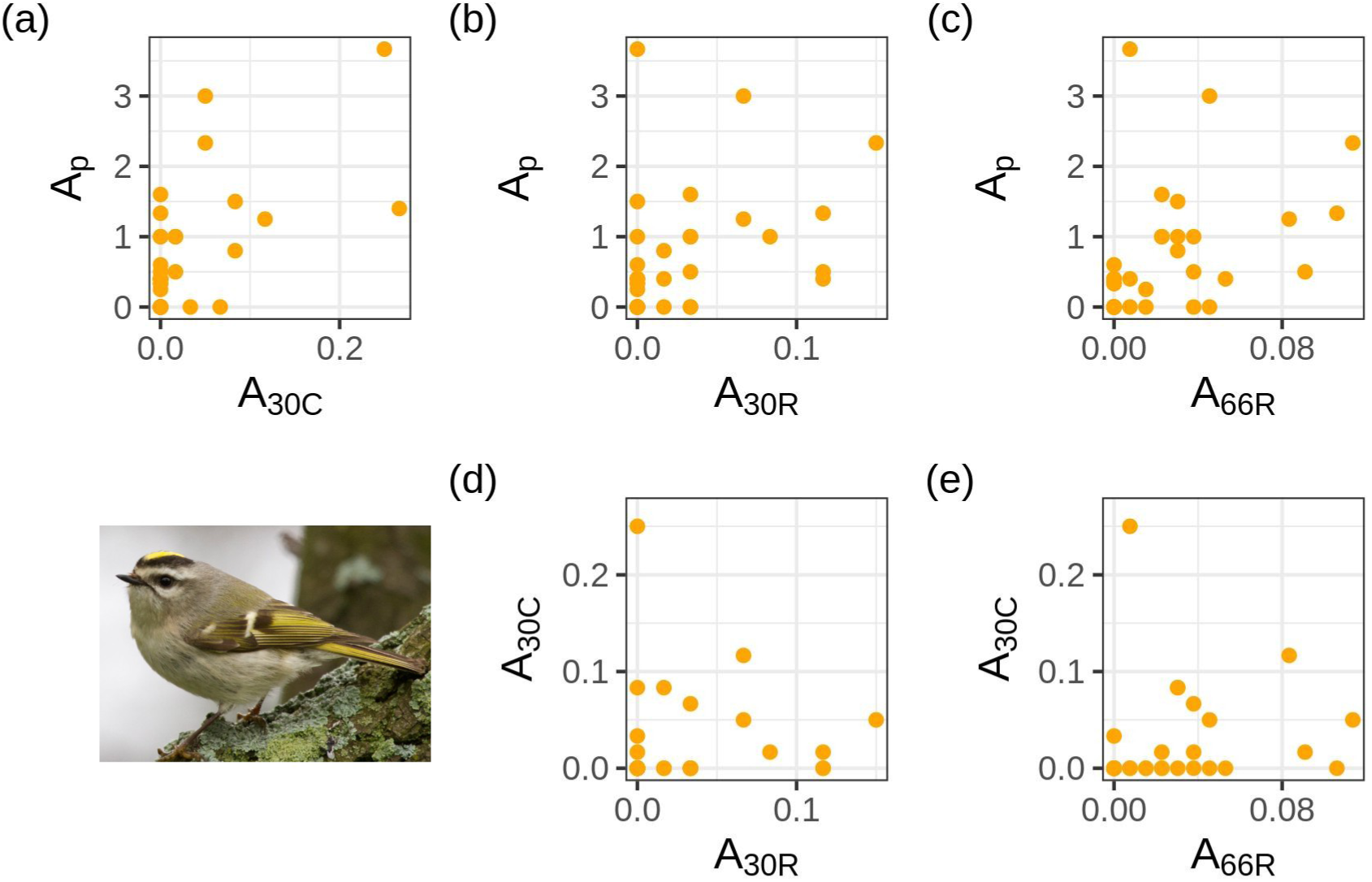
Correlation between daily observed values of relative abundance indices (Box 1) for Golden-crowned Kinglet. See Table 2 for Spearman’s correlation coefficients for each plot. Abundance indices for ARUs (the proportion of 30-second intervals with a vocalization) are correlated with the abundance index from point counts (mean number of individuals observed per count per day). Note that axis scales vary by abundance index; absolute values are less important here than the relationship between observations. Photo: “Golden-crowned Kinglet” by Laura Gooch, used under license CC BY-NC-SA 2.0. Cropped from original.

For Winter Wrens, observed abundance indices using randomly selected minutes from ARUs (*A*_30R,_ *A*_66R_) were more closely correlated with the observed abundance index from point counts (*A*_P_) than was the abundance index from 10 consecutive minute ARU surveys (*A*_30C_). For both species, *A*_P_ was most strongly correlated with *A*_66R_.

## 4 DISCUSSION

Importantly, abundance models trained with ARU data showed the increase in relative abundance indicating the arrival of migrants at the study site, suggesting that ARUs can be used to track migration phenology in stopover habitat for vocal species. ARUs also provided similar estimates of the number of species detected as point counts when analyzing randomly sampled minutes. Our results suggest that ARUs recording for an extended duration in migration stopover habitat can be just as effective as in-person point counts for monitoring migrating land birds.

Data from randomly selected minutes of ARU recordings detected more species and produced modeled abundance estimates that better showed the expected seasonal pattern of migration timing than data from consecutive minutes of ARU recordings. There are two likely explanations for this. First, randomly selected minutes are less temporally auto-correlated than consecutive minutes. For example, during a 10-minute in-person point count, little new information is gained during the seventh minute of the survey compared to what was collected during the sixth minute of the survey; a Winter Wren singing near the end of the sixth minute of a point count survey will likely still be singing in the beginning of the seventh minute. By selecting minutes randomly from across the five-hour survey window, the temporal correlation between each successive minute that is analyzed is minimized.

Second, during migration stopover, birds may move more and farther distances within the study area than they would during the breeding season, when they have established a territory. The community of birds within the immediate detection radius of an observer (either a person or a recording ARU) may therefore change over the course of five hours. Using randomly selected minutes provides a more complete sample of the birds using a spatial location over the entire course of the survey window.

For in-person point counts, the time taken to travel to a survey site takes up a major portion of the total time invested, so site visits are typically limited to once per day. With ARUs, no such constraints exist; it is possible to do multiple short-duration surveys from many locations over the course of one day without additional travel and field work logistics. We recommend that studies using ARUs on migration should randomly sample recordings of short periods of time (e.g. one-minute recordings) from a defined survey window relevant to the study question (e.g. the five hours following sunrise for passerines in temperate forest, or twilight to dawn for crepuscular and nocturnal species). In many studies using ARUs during the breeding season, researchers listened to consecutive minutes of recordings (e.g. 10 minutes or 2 minutes in Klingbeil & Willig (2015)). Given the improvement we saw when listening to random rather than consecutive minutes from an extended duration recording, we recommend that future studies using ARUs to monitor birds during wintering or breeding seasons test whether randomly selected minutes provide a more effective sample than consecutive minutes.

We did not detect an effect of either wind or rain in our model of the number of species detected. However, because we controlled for adverse weather conditions during our field surveys by not surveying on rainy or windy days, the number of high wind values in our data was low, as was the number of rainy survey days. We also noted anecdotally that occasionally the wind values recorded in person for a survey day did not correlate with the amount of wind heard while conducting our desk-based audio surveys; we speculate that wind direction in relation to the microphone may make a difference in how much wind is actually picked up by the ARU. Given that wind and rain have an effect on the detectability of birds in the study system (Ralph, Droege & Sauer, 1995), they remain important predictors to include, whether or not they appear significant in our model. Likewise, we suspect that the insignificance of the polynomial day of year term is due to sample size limitations.

Interpreting the significance of the noise variable is challenging, because we used the variable to describe all non-avian noise in the environment, which could include waves, airplanes, and frogs. We suspect that the overall significance of the noise variable may be due to frogs. Future studies may want to carefully consider whether they wish to distinguish between other vocalizing taxa and surrounding environmental noise. ARUs can be used to simultaneously sample multiple taxa (e.g. crickets and bats; Newson, Bas, Murray & Gillings, 2017) so researchers may want to incorporate analyses of non-bird biotic noise into study designs.

Estimates of abundance are more useful than estimates of occupancy for prioritizing conservation resources at dynamic temporal scales, such as during migration (Johnston et al., 2015). ARUs do not solve the problem of how to estimate true abundance in stopover habitat during migration. Imperfect detection means that the number of individuals detected is not necessarily a good estimate of the number of individuals present (MacKenzie & Kendall, 2002). Hierarchical models that account for imperfect detection (MacKenzie et al., 2002; Kéry & Royle, 2016) rely on assumptions about population closure that may be badly violated during migration, when birds do not adhere to territories and are present in stopover habitat for short periods of time. The period in which we can reasonably assume population closure for our study area during migration may be as short as several hours or as long as several days, depending on weather conditions. Therefore disentangling true occupancy or abundance from detectability is difficult, whether using traditional in-person survey methods or ARUs. Using a relative abundance index that does not account for detection probability could result in misleading estimates of abundance if detection probability is not constant (MacKenzie & Kendall, 2002). It is possible that individual birds’ vocalizations may increase over the spring migration period, as birds prepare for the breeding season. Using our abundance indices, increases in vocalizations would look like an increase in relative abundance, but the apparent increase would merely be an artifact of changing detectability. However, given the moderate (for Golden-crowned Kinglets) and strong (for Winter Wrens) correlation between *A*_p_ (observed abundance from point counts) and our ARU abundance indices, we believe increases in relative abundance seen in our model results (Fig. 3) are not mere artifacts of changes in detectability, but rather show real increases in abundance associated with the arrival of our focal species in the study area. ARUs can therefore provide valuable data about migration phenology and stopover habitat use, even if they cannot be used to estimate true abundance.

Differences in how well relative abundance models captured the arrival of migrants seemed to be partly dictated by how well aggregating detections by day reduced the number of “zero” counts in our data. For Golden-crowned Kinglets, the *A*_p_ and *A*_66R_ models seemed to be effective at detecting the increase in abundance associated with initial arrival, though they did not capture a peak of abundance, if one existed during the study period. It is unclear whether days with high counts of Golden-crowned Kinglets represent a real pulse of newly arrived birds, or the same birds already present in the region clustering more densely, or just random variation around a more or less constant number of individuals. A dense network of simultaneously recording ARUs in the region could help answer this question. Future studies might also consider increasing ARU survey effort beyond our maximum of 66 randomly selected minutes per day. The improvement in our models associated with increasing from 30 to 66 minutes suggests that increasing the number of minutes surveyed is beneficial.

Future studies using ARUs to monitor bird migration may wish to take advantage of ARUs’ unique ability to scale research in ways that may be infeasible or prohibitively expensive for in-person field work. For example, ARUs could be deployed in dense, small-scale networks to examine micro-habitat use in stopover regions. Alternatively, they could be deployed on a latitudinal gradient covering hundreds or thousands of kilometers to examine how vocal behavior changes over the spring migration period as birds approach their breeding grounds.

Applying the methods described here can facilitate an increase in survey effort in difficult-to-access migratory stopover habitat in high latitude forests. Temporal variation in accessibility in these habitats is dramatic, as unpaved roads typically turn from snow to slush to impassable mud before hardening into reliably dry surfaces in early summer. ARUs can eliminate many of the restrictive logistics and safety concerns for researchers interested in monitoring spring migration. Our method of using desk-based surveys of randomly selected minutes from ARUs can be used by any researcher with the skills to conduct point counts. Researchers can set up ARUs during winter conditions when access to study sites over snow is relatively easy (e.g. using snowmobiles, skis or snowshoes), and revisit to collect the audio data once conditions have stabilized in late spring. Our methods for using ARU data to model relative abundance of focal species and the number of species present during migration can be immediately applied to increase monitoring effort in logistically difficult regions.

## Acknowledgments

This work was funded by grants to ER from the Wilson Ornithological Society and the Copper Country Audubon Society. Land access and logistical support was provided by the Keweenaw Land Trust and Pat Toczydlowski. We are deeply grateful to Calvin and Steve Koski, Mark Summersett, and Bob and Nancy Korth for their support and assistance with transportation to the study site in difficult and unpredictable conditions. Our thanks also to Joseph Youngman, David Flaspohler, Drew Meyer and Dana Neufield for assistance with the pilot year surveys, and to Jon Yearsley, Hannah White, and the Ecological Modeling lab at University College Dublin for consultation on modeling methods. WG was funded during this work by Science Foundation Ireland grant number 15/IA/2881.

## Author Contributions

ER and WG developed the study design and methodology; ER prepared for and performed all field work, and conducted all desk-based audio surveys; ER and WG conducted the data analysis, wrote the manuscript, and gave final approval for publication.

## Data Availability Statement

Data and code to reproduce analyses can be found at https://doi.org/10.5281/zenodo.3964500 (ellieroark, 2020). Audio recording files used to produce this analysis are archived at https://doi.org/10.5281/zenodo.3964574 (Roark & Gaul, 2020).

## Notes

https://doi.org/10.5281/zenodo.3964500

https://doi.org/10.5281/zenodo.3964574

### Competing Interest Statement

The authors have declared no competing interest.

