## Appendix A and Supplementary Tables and Figures for "Monitoring migratory birds in stopover habitat: assessing the value of extended duration audio recording"

### Appendix A: Supplementary Methods and Results

### METHODS

#### Pilot year surveys; site selection; date selection

We selected survey sites randomly across the study area with the criteria that points were at least 100 m away from the shoreline, and at least 300 m apart, to ensure ARUs recorded non-overlapping areas (Klingbeil & Willig, 2015). During the pilot year of surveys in 2018, nineteen points were initially selected and tested (eighteen chosen randomly as previously mentioned, and one point selected by hand near the tip of the peninsula). Three of those initial nineteen points were dropped for the 2019 season, due to difficult access, posted private property signs, and smaller effective survey area (because of proximity to water) relative to the other points. For the 2019 field season, the survey area was expanded from 2.5 $km^2$ to 2.7 $km^2$ to include an adjacent Keweenaw Land Trust property, and two additional survey points were added.

We selected survey dates by reviewing historic observations for four early season migrant species: Golden-crowned Kinglet (*Regulus satrapa*), Northern Flicker (*Colaptes auratus*), Winter Wren (*Troglodytes hiemalis*), and Hermit Thrush (*Catharus guttatis*). For each of these species, we downloaded all Michigan eBird records from 1970 to 2018 (eBird Basic Dataset, 2018), and used the number of daily detections (without attempting to correct for variable survey effort or detectability) to select field work dates that would give us a good chance of surveying through the early migration period (i.e. starting before any individuals of the focal species were present and continuing through a period of high focal species abundance).

To determine our survey period for the 2019 field season, we subset eBird data for the region to observations made between 1 January and approximately 8 July (day of year 190), in the four counties nearest our survey area (Baraga, Keweenaw, Houghton, Marquette). We plotted day of year by the number of observations per species (as a proxy for abundance) to determine both the day of arrival at the study area and the approximate peak of migration for each species. We selected 1$^{st}$ April (day of year 91) as the start date for surveys because in eBird records from previous years, 90% of Golden-crowned Kinglet observations and 99% of records for the other three target species occurred after this date. We wanted to balance the benefit of being early enough to catch the first migrants against the cost of potentially spending weeks in a remote field location without our focal species present. Plots of cumulative observations by day of year showed that surveying through day of year 141 (May 22) would allow us to catch the peak in the daily number of observations reported to eBird for each of our target species. This survey period did not encompass the entirety of spring migration for all migrant species in the region, but was designed to capture the peak of abundance during migration for our focal migrant species.

**Point count protocols**

The field technician announced aloud the beginning and end of each point count, as well as the date, time, location name, and geographic coordinates, so that this information could be recorded on the ARU, as well as on the technician's data sheet. Each species observed was noted on a data sheet, including the number of individuals seen, the bearing of the first individual or group detected (relative to the direction the observer was facing), the detection method (call, song, woodpecker drum or visual), the distance from the observer (in three distance bands of 0–25 m, 26–50 m, or 50+ m), and the minute of the survey in which the species was first detected (0–9). The observer also noted cloud cover (0–33%, 34–66%, 67–100%), precipitation (Dry, Fog/Haze, Drizzle, Rain/Snow), Beaufort wind scale rating (0-5) (Beaufort, 1805), and non-bird noise level for each point count (0–4). We did not survey if the wind was greater than force 5, or in heavy, continuous precipitation.

**Desk-based survey protocols**

No more than five hours of desk-based audio surveys were conducted in a single day, and all audio recordings were listened to at full speed. The technician was allowed a maximum of 15 minutes to listen to each 10 minute recording, during which time they could pause, rewind or replay the audio file, and could look up and play songs or calls from any external resource they felt may be helpful, excluding using any kind of automated identification program. Because of the difficulty of deciding what constitutes a single "vocalization" from species with different songs and calls, we did not attempt to count the number of vocalizations. While conducting desk-based audio surveys, we viewed spectrograms of the recording in Audacity (Audacity Team, 2019). Spectrograms were viewed in gray scale, with a minimum frequency of 0 kHz and a maximum frequency of 15 kHz. The gain (brightness) of the spectrogram was 20 dB, while the range (contrast) was 80 dB. Frequency gain was 0 dB/dec. Window size was 256, with window type Hanning and a zero padding factor of 1.

To process the data from the 10 consecutive minute counts, we clipped the audio recording of each 10 minute point count from its larger audio file, excluding voice announcements about the location, date and time of recording and including only the "begin point count" and "end point count" announcements from the survey technician. Because the field survey technician (ER) also conducted the desk-based audio bird surveys, the second author (WG) anonymized the audio recorder file names so that ER could not see or hear the dates and locations of the audio recordings. This reduced the possibility that memories of particular days or locations would influence the data collected during the desk-based audio survey. In the anonymized file names we included an indicator of the two week period in which the point count took place ("early" or "late" in April or May) because information about season is used by bird observers to inform their mental list of “possible” species, and this information would be available to a technician conducting desk-based audio bird surveys in practical applications.

In order to ensure that the technician’s desk-based audio survey species identifications were reproducible, we duplicated 20% of the 10 consecutive minute recordings, and assigned new anonymous names to the duplicated recordings, so that the technician listened to that data twice. After data entry and de-anonymization, we compared the species detected in each duplicated recording.

The desk-based survey process used for listening to 24 random minutes was similar to the process used for 10 consecutive minute recordings. Selection of the 24 random minutes was done in R using the ‘warbleR’ package and work flow (Araya-Salas & Smith-Vidaurre, 2017; R Core Team, 2020). We wanted to analyze a minimum of 20 random minutes without anthropogenic disturbance, so we selected and clipped 24 minute-long segments from each day’s audio for each recorder. We subsequently discarded any clip that contained a human voice, but did not discard clips containing other possibly anthropogenic noise (e.g. footsteps), because we did not have a clear way to distinguish between human and wild animal sounds. We also did not control for distant anthropogenic noises such as vehicles and planes. We created a new sample of 24 random minutes to select from each ARU on each survey day, so that we did not select the same minutes from each day or ARU. We only selected 24 random minutes from an ARU on days when the unit recorded the full five-hour survey window. We listened to randomly selected minutes using the procedures outlined above for the 10 consecutive minute desk-based audio surveys, but allowing for a maximum of 50 minutes to listen to each set of 24 random minutes. This allowed for approximately the same effective listening time for the audio files (50% more than the length of the original file), but included extra time for file management (opening and closing the audio files in Audacity).

**Sample size on 16th and 17th April**

The number of surveys per day over the course of the season varied based on local conditions. We deployed at least three ARUs every day, and deployed a fourth ARU on days when insufficient weatherproofing was not likely to interfere with recording efforts. We considered all survey days with at least three in person point counts, and five hour recording windows from at least three ARUs, a "complete" survey day. On two days (April 16th and 17th) we failed to capture a complete survey day due to ARU SWIFT03 malfunctioning. On April 16th, we also were unable to conduct three in-person point counts, and conducted only two counts, one alongside the functionally recording SWIFT01, and one next to the malfunctioning SWIFT03. SWIFT02 successfully recorded the full five hour survey window on April 16th, but no point count was conducted there.

Because these dates coincided with an important arrival period of migrants into the study area, we did not drop them from our analyses. Because we modeled species richness per count rather than per day, those models were unaffected by the anomalies described above. However, because we aggregated our abundance indices per day, it is important to note that the abundance indices for April 16th and 17th are different than the other survey days (see Box 1 for description of abundance indices). On the 16th of April, the abundance index for point counts is the mean number of individuals detected per count, but averaged between only two point counts instead of the usual three counts. On that date, the abundance index for consecutive minute ARU counts was $A_{10C}$, rather than $A_{30C}$. On April 17th, the abundance index for consecutive minute ARU counts was $A_{20C}$, rather than $A_{30C}$. We opted to leave these dates in our models with reduced survey effort, rather than remove them. To compensate for the malfunctioning third recorder, we selected an additional 11 random minutes from each of the functional recorders so that we do have abundance indices of $A_{66R}$ and $A_{30R}$ for those dates, but they are sampled from only two ARUs instead of three ARUs.

For days when four ARUs were deployed, we listened to 22 random minutes from each ARU (88 minutes total), but standardized effort to 66 minutes and 30 minutes per day for the $A_{66R}$ and $A_{30R}$ indices, respectively. To do this, we randomly selected 66 and 30 minutes from the entire survey day, which may include data from all four of the ARUs.

**Generalized Linear Mixed Model for the number of species detected**

We included an interaction between day of year and survey type because we anticipated that the effect of survey type might change over time, for example if increased numbers of species or individuals later in migration made distinguishing identifiable sounds on the audio recordings more difficult. We included an interaction between rain and survey type because we expected that even small amounts of rain hitting the ARUs might impair our ability to detect birds on desk-based audio surveys more than a similar amount of precipitation during point counts. Because the number of observations for some values of the categorical weather variables was small, we pooled levels as follows: wind was binned into categories 0–1, 2, or 3+; rain was binned into “wet” and “dry” conditions; noise was binned as 0, 1, or $\geq$2. We centered and scaled all continuous variables. We did not perform model selection but rather included all variables in the final model due to an *a priori* expectation that all variables were relevant to the study system. We assigned the weather variables noted in person during point counts to all the desk-based audio surveys from the same unit on the same day. While weather conditions may have changed slightly over the course of the survey window, the observed weather conditions from the point count represent our best estimate of the conditions at each survey location on each survey day.

**Boosted Regression Tree (BRT) relative abundance models**

We fit BRTs with a Laplace distribution, and an absolute loss link function which is more robust than a RMSE loss function to data with long tailed distributions (Hastie, Tibshirani, & Friedman, 2009). We ran BRTs with an interaction depth of one, a minimum of one observation per node, and a bag fraction of 0.8. To optimize the number of trees and shrinkage parameter for our boosted regression tree models (BRTs), we set the cross validation parameter built into the gbm function to ten, and looked at graphs of the cross-validation test error for ten iterations of our model using the function gbm.perf in the gbm package (Greenwell, Boehmke, Cunningham and GBM Developers, 2019; R Core Team, 2020). We aimed to optimize the shrinkage parameter to grow at least 1000 trees before models started to overfit (indicated by increasing cross-validation test error) (Elith, Leathwick & Hastie, 2008). We tested shrinkage parameter values ranging from 0.01 to 0.0001, and graphed test error when adding up to 10,000 trees (Figs S4-S5). We chose our final shrinkage parameter and number of trees for each model based on visually assessing plots of cross-validation test error, attempting to avoid over fitting as much as possible for each particular model iteration (Figs S4-S5). We opted to use a single shrinkage parameter and number of trees for all iterations of each relative abundance model, rather than attempting to automatically tune the shrinkage parameter and number of trees within each iteration. Because our final predictions were the averages from 1000 model iterations, our results should not be unduly impacted even if individual model iterations did not achieve the minimum possible test error. We ultimately chose the following shrinkage parameters and number of trees for each of our models: shrinkage parameter of 0.0005 and 3000 trees for Winter Wren models $A_p$, $A_{30R,}$ $A_{66R}$; shrinkage parameter of 0.0001 and 3000 trees for Winter Wren model $A_{30C}$ and Golden-crowned Kinglet models $A_{30C}$ and $A_{30R}$; shrinkage parameter of 0.0005 and 2000 trees for Golden-crowned Kinglet models $A_p$ and $A_{66R}$.

In some cross-validation folds, the test error increased immediately, after fitting only one tree (Fig. S5 b). In these cases, we tested the smallest possible shrinkage parameter recommended by Elith, Leathwick & Hastie (2008), which was 0.0001. We believe that the immediate over fitting is likely due to the large number of zeros in the data, and we think it unlikely that we would be able to tune parameters to build good models with these data; the problem is with the data rather than with the tuning of the model parameters (see e.g. the large number of observed zeros in Fig. 3 b, f). This supports our overall conclusion that using consecutive minute ARU recordings is less useful for assessing the relative abundance of migrant species than using randomly selected minutes from ARU recordings.

To ensure that our results were not solely based on our choice of modeling method, we also modeled relative abundance using generalized additive models (GAMs) because they allowed specification of a negative binomial error distribution which we suspected might fit our data well. We tested GAMs using two of our four abundance indices: $A_p$ and $A_{30C}$. GAMs were fit with a thin plate spline, with the number of knots optimized at k = -1. GAMs were fit using the ‘gam’ function in the ‘mgcv’ package (Wood, 2003; Wood, 2011; Wood, 2017). As with BRTs, we fit GAMs using 200 iterations of five-fold temporal block cross validation, that used blocks of three consecutive days (Fig. S3). To compare GAMs and BRTs, we calculated a mean Root Mean Square Error (RMSE) of all 1000 model iterations.

### RESULTS

#### Analysis of duplicated recordings

The duplicated recordings had perfect agreement about the occurrence of Winter Wren, indicating that we can have high confidence in Winter Wren identification from ARU surveys. We intended to use Krippendorff’s alpha (Krippendorff, 2013) to assess agreement about detections of focal species on duplicated desk-based audio surveys of 10 consecutive minute counts. However, we did not have enough detections of Golden-crowned Kinglet in our duplicated recordings to calculate a value for Krippendorff’s alpha; we therefore do not have an estimate of the reliability of identification of Kinglets on ARU recordings. We encourage future researchers to give consideration to listener agreement when using ARU data.

### GAM Results

GAMs and BRTs performed similarly for estimating relative abundance trends across both survey types on which GAMs were tested based on evaluating Root Mean Square Error (RMSE) (Table S2), and gave qualitatively similar models of the change in bird abundance during the migration season. This indicated that the choice of modeling method and error distribution did not have a large effect on the results of our model, and we ultimately chose BRTs following Johnston et al.'s (2015) methods for modeling abundance.

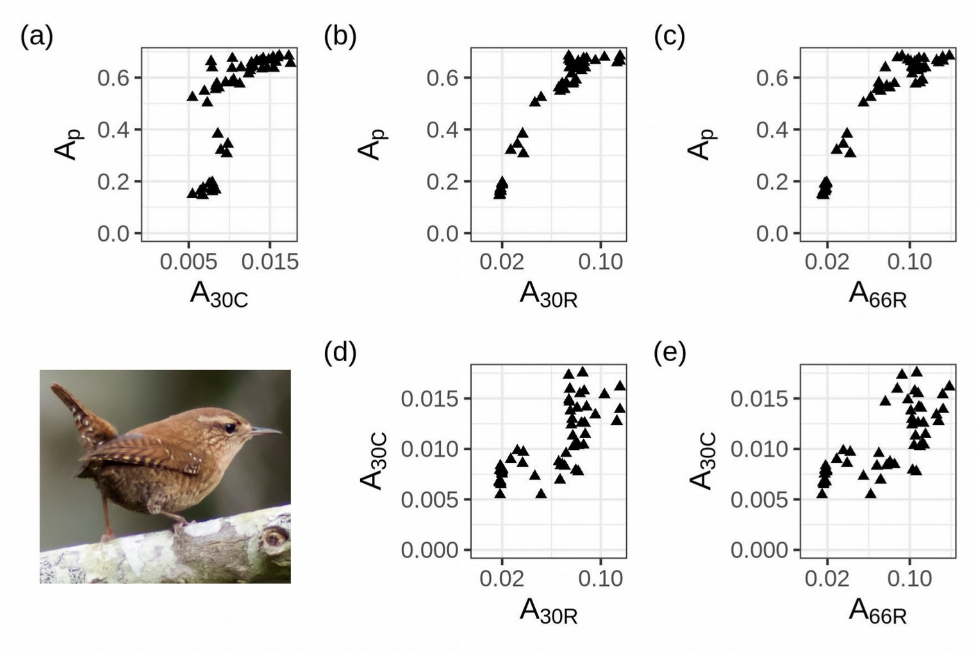


**Figure S1:** Correlation between predicted relative abundance index values for Winter Wren. Predictions are the mean calculated from 1000 iterations of a Boosted Regression Tree model. Spearman's rank correlation coefficients can be found in Table 2. Predictions of index values are strongly correlated, indicating that the model is finding the same signal regardless of whether training data are from ARUs or point counts. Note that axis scales vary by abundance index; absolute values are less important here than the relationship between predictions. Photo: "Winter Wren" by ilouque, used under license CC BY 2.0. Cropped from original.

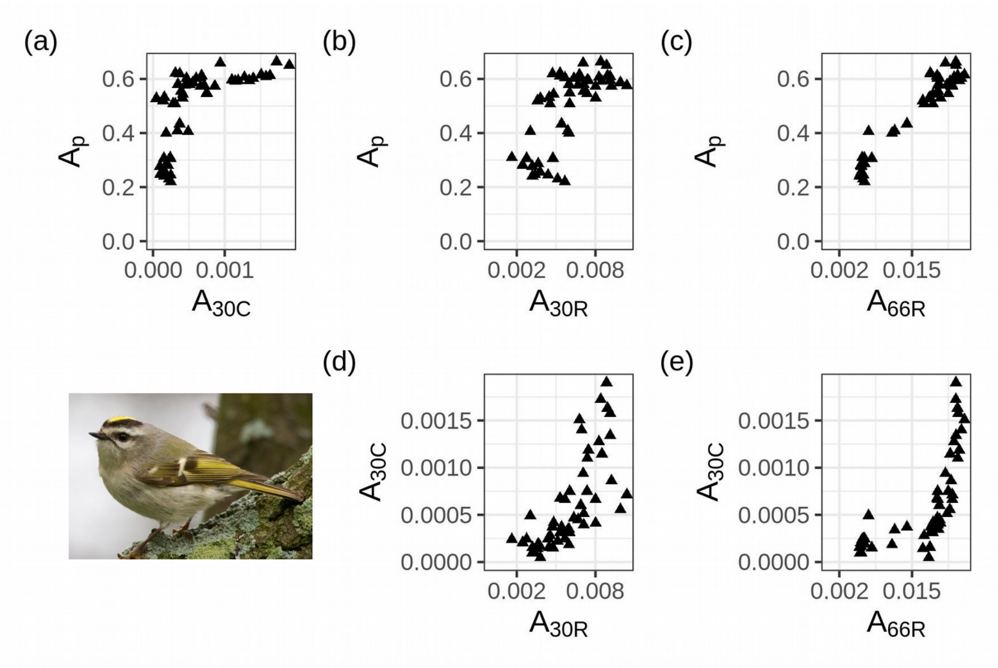


**Figure S2:** Correlation between predicted relative abundance index values for Golden-crowned Kinglet. Predictions are the mean calculated from 1000 iterations of a Boosted Regression Tree model. Spearman's rank correlation coefficients can be found in Table 2. Predictions of index values are strongly correlated, indicating that the model is finding the same signal regardless of whether training data are from ARUs or point counts. Photo: "Golden-crowned Kinglet" by Laura Gooch, used under license CC BY-NC-SA 2.0. Cropped from original.

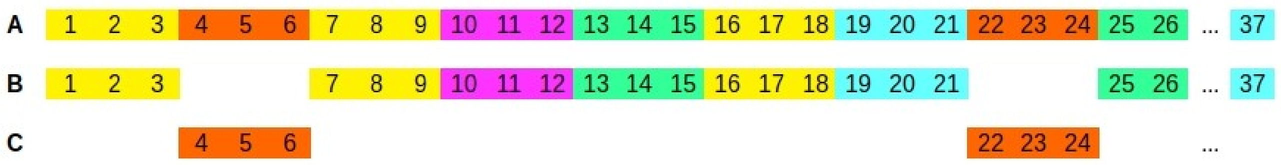


**Figure S3:** Schematic of temporal block cross-validation used for fitting and testing abundance models. Days (1, 2, 3, ..., 37) were grouped into blocks of three consecutive days. Each block of three days was assigned to one of five cross-validation folds (colors, panel A). Abundance models were fitted by withholding data from days in one fold (e.g. the orange fold) and using data from days in the other four folds as training data (B). The performance of the model was evaluated based on how well it predicted data from days in the test fold (C).

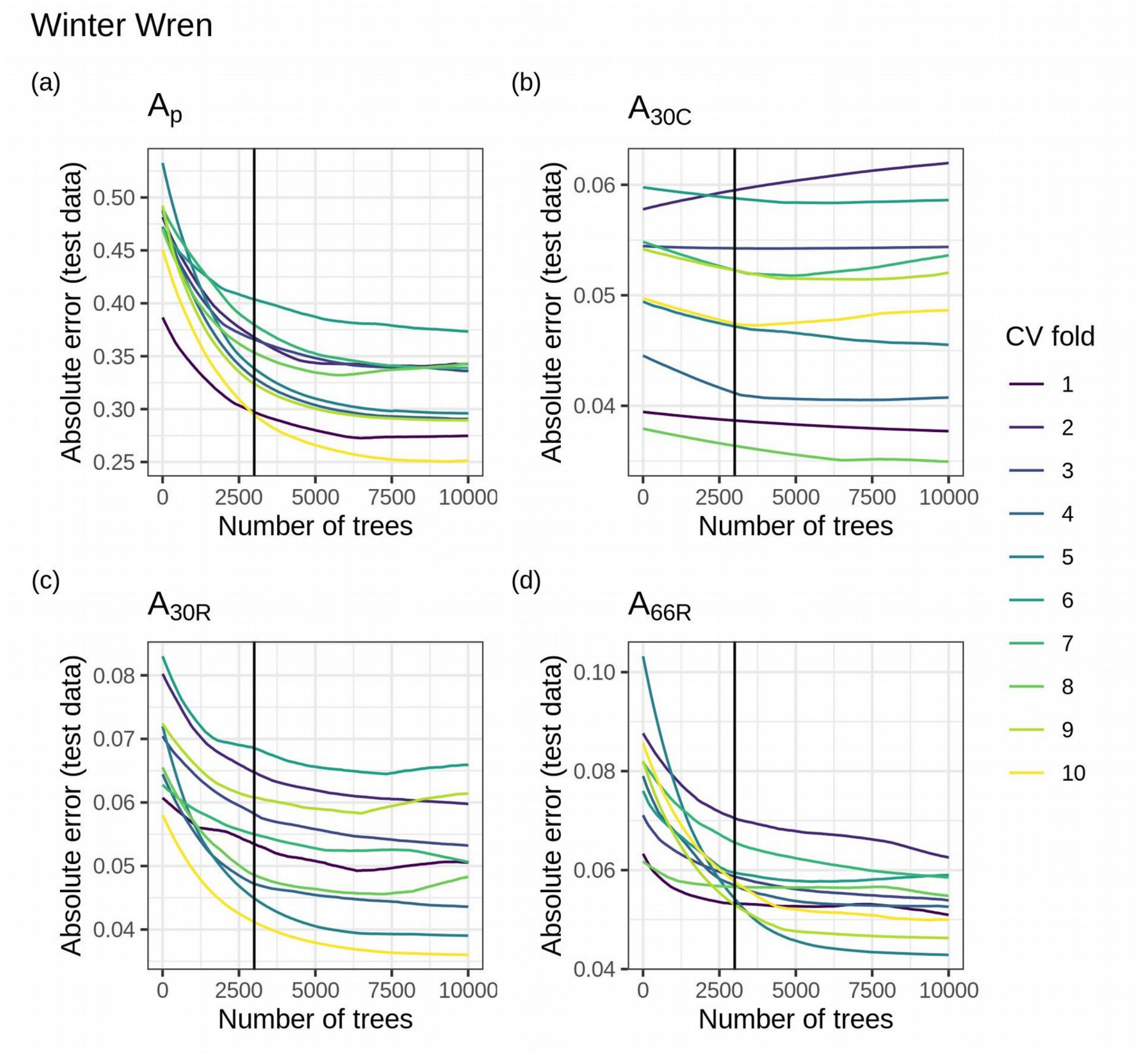


**Figure S4:** Test error for 10 cross validation folds for a Boosted Regression Tree model, for each of four abundance indices for Winter Wren relative abundance models. Colored lines show the cross-validation fold for which the error was calculated, and the black vertical bar shows the number of trees we chose for each abundance index. Plots a, c and d all show test error for a shrinkage rate of 0.0005; plot b shows test error for a shrinkage rate of 0.0001. For models that appeared to overfit immediately (b), we believe the problem was with the data rather than with the tuning parameters (see discussion in Appendix A).

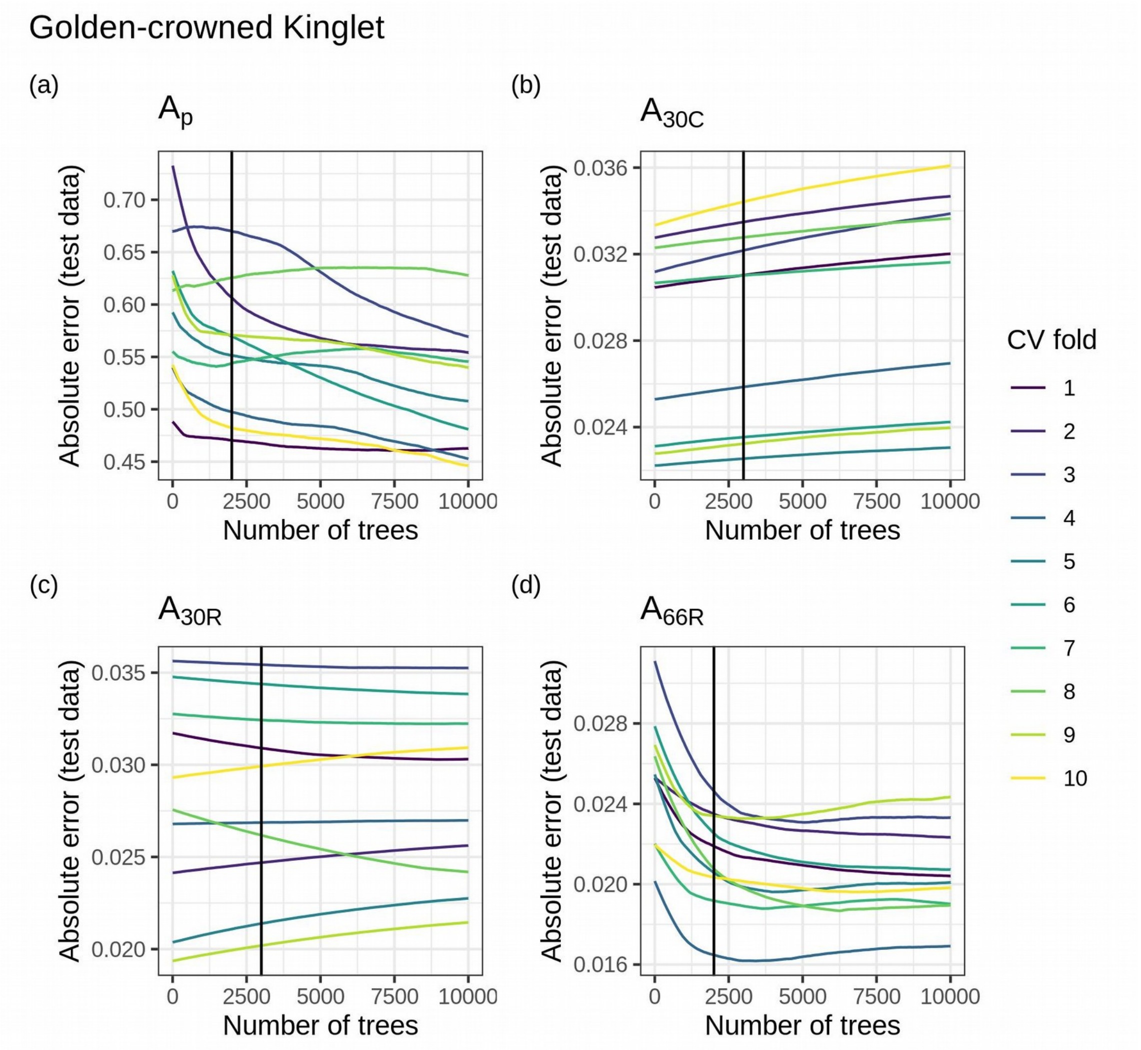


**Figure S5:** Test error for 10 cross validation folds for a Boosted Regression Tree model, for each of four abundance indices for Golden-crowned Kinglet relative abundance models. Colored lines show the cross-validation fold for which the error was calculated, and the black vertical bar shows the number of trees we chose for each abundance index. Plots a and d all show test error for a shrinkage rate of 0.0005; plots b and c show test error for a shrinkage rate of 0.0001. For models that appeared to overfit immediately (b, c), we believe the problem was with the data rather than with the tuning parameters (see discussion in Appendix A).

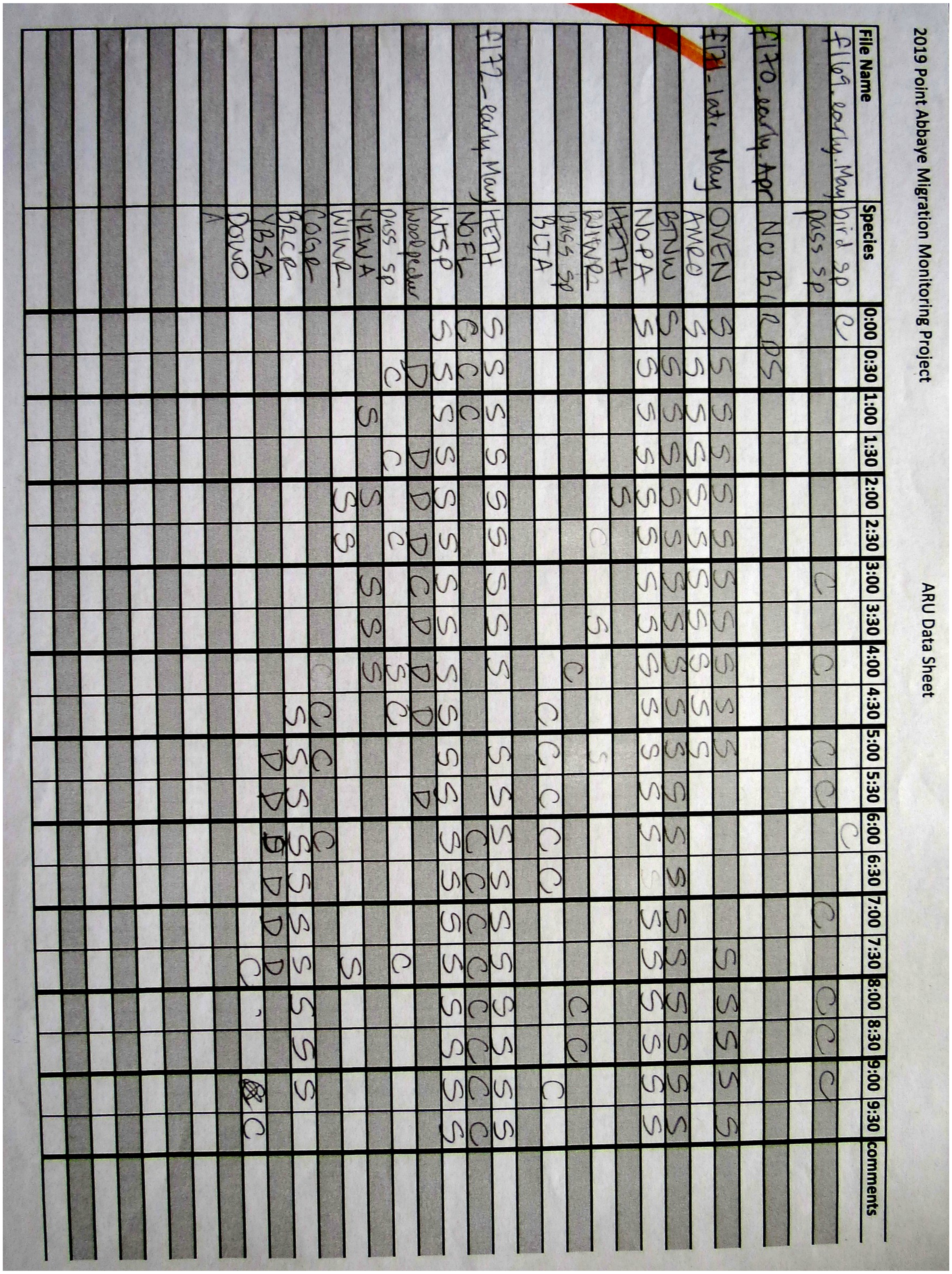


**Figure S6:** Example data sheet for desk-based surveys of 10 consecutive minute ARU recordings. Species codes are 4-letter alpha codes; see Appendix A, “Desk-based survey protocols” for details about vocalization codes recorded in each 30-second interval.

**Table S1:** Species detected during the 2019 field season on the Point Abbaye peninsula, showing frequency of point count detection type for each species, and indicating whether species were detected with ARUs. *denotes species detected only using ARUs.

| Scientific Name | Common Name | Point Count Detections | | | | ARU Detection? |
|---|---|---|---|---|---|---|
| | | Visual | Call | Song | Drum | |
| *Branta canadensis* | Canada Goose | 0 | 5 | 1 | 0 | yes |
| *Anas platyrhynchos* | Mallard | 0 | 0 | 0 | 0 | yes* |
| *Meleagris gallopavo* | Wild Turkey | 0 | 0 | 0 | 0 | yes* |
| *Antigone canadensis* | Sandhill Crane | 0 | 1 | 0 | 0 | no |
| *Gavia immer* | Common Loon | 0 | 2 | 0 | 0 | no |
| *Cathartes aura* | Turkey Vulture | 2 | 0 | 0 | 0 | no |
| *Haliaeetus leucocephalus* | Bald Eagle | 4 | 7 | 0 | 0 | yes |
| *Sphyrapicus varius* | Yellow-bellied Sapsucker | 2 | 14 | 0 | 21 | yes |
| *Dryobates pubescens* | Downy Woodpecker | 6 | 5 | 1 | 4 | yes |
| *Dryobates villosus* | Hairy Woodpecker | 5 | 5 | 0 | 13 | yes |
| *Colaptes auratus* | Northern Flicker | 0 | 25 | 0 | 0 | yes |
| *Dryocopus pileatus* | Pileated Woodpecker | 1 | 6 | 0 | 4 | yes |
| *Falco sparverius* | American Kestrel | 1 | 0 | 0 | 0 | no |
| *Falco columbarius* | Merlin | 0 | 2 | 0 | 0 | yes |
| *Empidonax flaviventris* | Yellow-bellied Flycatcher | 0 | 0 | 1 | 0 | no |
| *Empidonax minimus* | Least Flycatcher | 0 | 0 | 4 | 0 | yes |
| *Sayornis phoebe* | Eastern Phoebe | 1 | 0 | 0 | 0 | no |
| *Vireo solitarius* | Blue-headed Vireo | 0 | 0 | 10 | 0 | yes |
| *Cyanocitta cristata* | Blue Jay | 0 | 19 | 0 | 0 | yes |
| *Corvus brachyrhynchos* | American Crow | 1 | 12 | 0 | 0 | yes |
| *Corvus corax* | Common Raven | 0 | 28 | 0 | 0 | yes |
| *Poecile atricapillus* | Black-capped Chickadee | 0 | 38 | 23 | 0 | yes |
| *Sitta canadensis* | Red-breasted Nuthatch | 0 | 3 | 23 | 0 | yes |
| *Sitta carolinensis* | White-breasted Nuthatch | 0 | 3 | 8 | 0 | yes |
| *Certhia americana* | Brown Creeper | 3 | 18 | 24 | 0 | yes |
| *Troglodytes hiemalis* | Winter Wren | 0 | 0 | 62 | 0 | yes |
| *Regulus satrapa* | Golden-crowned Kinglet | 1 | 34 | 10 | 0 | yes |
| *Regulus calendula* | Ruby-crowned Kinglet | 6 | 0 | 18 | 0 | yes |
| *Catharus minimus* | Gray-cheeked Thrush | 1 | 0 | 0 | 0 | no |
| *Catharus ustulatus* | Swainson's Thrush | 2 | 0 | 1 | 0 | yes |
| *Catharus guttatus* | Hermit Thrush | 6 | 10 | 9 | 0 | yes |
| *Turdus migratorius* | American Robin | 2 | 22 | 14 | 0 | yes |
| *Toxostoma rufum* | Brown Thrasher | 0 | 0 | 0 | 0 | yes* |
| *Pinicola enucleator* | Pine Grosbeak | 0 | 1 | 0 | 0 | no |
| *Haemorhous purpureus* | Purple Finch | 0 | 0 | 10 | 0 | yes |
| *Spinus pinus* | Pine Siskin | 0 | 1 | 0 | 0 | no |
| *Spinus tristis* | American Goldfinch | 0 | 13 | 1 | 0 | yes |
| *Spizelloides arborea* | American Tree Sparrow | 1 | 0 | 0 | 0 | no |
| *Spizella passerina* | Chipping Sparrow | 1 | 0 | 1 | 0 | no |
| *Passerella iliaca* | Fox Sparrow | 0 | 0 | 1 | 0 | no |
| *Melospiza melodia* | Song Sparrow | 0 | 0 | 1 | 0 | no |
| *Zonotrichia albicollis* | White-throated Sparrow | 2 | 4 | 26 | 0 | yes |
| *Junco hyemalis* | Dark-eyed Junco | 0 | 6 | 3 | 0 | yes |
| *Molothrus ater* | Brown-headed Cowbird | 0 | 0 | 1 | 0 | no |
| *Quiscalus quiscula* | Common Grackle | 1 | 11 | 0 | 0 | yes |
| *Seiurus aurocapilla* | Ovenbird | 1 | 0 | 11 | 0 | yes |
| *Parkesia noveboracensis* | Northern Waterthrush | 0 | 0 | 1 | 0 | yes |
| *Mniotilta varia* | Black-and-white Warbler | 1 | 1 | 6 | 0 | yes |
| *Oreothlypis ruficapilla* | Nashville Warbler | 0 | 0 | 1 | 0 | yes |
| *Setophaga ruticilla* | American Redstart | 0 | 0 | 7 | 0 | yes |

| Scientific Name | Common Name | Point Count Detections | | | | ARU Detection? |
|---|---|---|---|---|---|---|
| | | Visual | Call | Song | Drum | |
| *Setophaga tigrina* | Cape May Warbler | 0 | 0 | 0 | 0 | yes* |
| *Setophaga americana* | Northern Parula | 0 | 0 | 12 | 0 | yes |
| *Setophaga fusca* | Blackburnian Warbler | 0 | 0 | 0 | 0 | yes* |
| *Setophaga petechia* | Yellow Warbler | 0 | 0 | 2 | 0 | no |
| *Setophaga caerulescens* | Black-throated Blue Warbler | 0 | 0 | 0 | 0 | yes* |
| *Setophaga palmarum* | Palm Warbler | 1 | 0 | 0 | 0 | no |
| *Setophaga pinus* | Pine Warbler | 0 | 0 | 1 | 0 | yes |
| *Setophaga coronata* | Yellow-rumped Warbler | 2 | 2 | 47 | 0 | yes |
| *Setophaga virens* | Black-throated Green Warble | 0 | 0 | 28 | 0 | yes |
| *Pheucticus ludovicianus* | Rose-breasted Grosbeak | 0 | 0 | 0 | 0 | yes* |
| NA | passerine sp. | 3 | 14 | 7 | 0 | yes |
| NA | woodpecker sp. | 0 | 3 | 0 | 17 | yes |
| NA | bird sp. | 0 | 31 | 0 | 0 | yes |
| NA | duck sp. | 1 | 0 | 0 | 0 | no |
| NA | gull sp. | 0 | 1 | 0 | 0 | no |
| NA | warbler sp. | 1 | 1 | 0 | 0 | yes |
| NA | vireo sp. | 0 | 0 | 1 | 0 | no |
| NA | corvid sp. | 0 | 0 | 0 | 0 | yes |
| NA | flycatcher sp. | 0 | 0 | 0 | 0 | yes |

**Table S2:** Root Mean Square Error (RMSE) for Boosted Regression Trees (BRT) and Generalized Additive Models (GAM) of relative abundance for focal species. See Box 1 for abundance index definitons.

| Abundance Index | Winter Wren | | Golden-crowned Kinglet | |
|---|---|---|---|---|
| | BRT | GAM | BRT | GAM |
| $A_p$ | 0.466 | 0.545 | 0.781 | 0.771 |
| $A_{30C}$ | 0.079 | 0.077 | 0.057 | 0.056 |
